# Distinct spatial associations of adversity with hippocampal macro- and microstructure in early adolescence

**DOI:** 10.64898/2026.07.30.741706

**Authors:** Mylla Marsiglia, Doruk Yigit Erigüc, Bianca Serio, Meike D Hettwer, Jordan DeKraker, Laura Waite, Felix Hoffstaedter, Boris C Bernhardt, Simon B Eickhoff, Sofie L Valk

## Abstract

Youth adversity has been associated with alterations in hippocampal structure; however, it remains unclear whether different forms of adversity relate distinctly to its organization across the anterior-posterior and proximal-distal axes. Here, we investigated the associations of different types of adversity with multiple structural properties of the hippocampus in 5,263 early adolescents from the ABCD Study. Hippocampal macrostructure was characterized using volume, thickness, and gyrification, whereas T1w/T2w ratio served as an *in vivo* proxy for microstructure. Adversity was assessed at the family level using questionnaires on family environment and parenting, and at the socioeconomic level using income-to-needs ratio and neighborhood disadvantage measured by the Area Deprivation Index. Associations were examined for each adversity type separately in multi-variate analyses and for cumulative adversity exposure in univariate models. Hippocampal features were obtained using HippUnfold, an advanced automatic segmentation approach that accounts for interindividual folding variability, and were analyzed globally as well as across the hippocampal anterior-posterior and proximal-distal axes. Socioeconomic measures showed widespread associations with hippocampal macrostructure across the whole hippocampus and both anatomical axes, whereas associations with T1w/T2w ratio were limited and observed only for neighborhood disadvantage along the anterior-posterior axis. Cumulative adversity exposure was consistently associated with alterations in CA1 and subiculum across volume, thickness, and gyrification, but not T1w/T2w ratio. Together, these findings suggest that different types of adversity exhibit distinct spatial associations across complementary hippocampal macro- and microstructural features, highlighting regional variation in the susceptibility of the developing hippocampus to environmental adversity.

## Introduction

The hippocampus is a highly plastic structure that undergoes a protracted maturational period during adolescence (Langnes et al., 2020; Tamnes et al., 2018; Zatorre et al., 2012). This developmental window is characterized by heightened plasticity, during which neural circuits remain especially sensitive to environmental influences, potentially increasing susceptibility to adverse experiences (Fox et al., 2010; Knudsen, 2004; Luby et al., 2020; McLaughlin et al., 2019; Sisk and Gee, 2022; Tooley et al., 2021). Adversity may influence hippocampal development through stress-response systems, including the activation of the hypothalamic-pituitary-adrenal (HPA) axis and downstream glucocorticoid signaling (Gunnar and Quevedo, 2007). The hippocampus has been extensively described as a region susceptible to stress due to its high density of glucocorticoid receptors (Christian et al., 2011; Gerlach and McEwen, 1972; Liu et al., 1997; McEwen and Sapolsky, 1995; McEwen, 1999; Wang et al., 2013). Activation of these receptors can initiate molecular processes that lead to structural remodeling of the hippocampus, such as neurogenesis suppression and dendritic atrophy (Christian et al., 2011; McEwen, 1999).

The anatomical organization of the hippocampus comprises multiple subfields that can be characterized along two principal axes. Along its transverse (proximal-distal) axis, the hippocampus extends from the subicular complex, which lies adjacent to the neocortex at the proximal end, through the cornu ammonis (CA1-4) regions, to the dentate gyrus (DG) at the distal end (Amunts et al., 2005). Its long (anterior-posterior) axis spans from the posterior (tail) to the anterior (head) portion (Nichols et al., 2023; Plachti et al., 2019; Poppenk et al., 2013). Both axes reflect differences in functional specialization, cytoarchitecture, connectivity, and maturational patterns across development (Amunts et al., 2005; Karat et al., 2023; Langnes et al., 2020; Morris et al., 2024; Nichols et al., 2023; Plachti et al., 2019; Poppenk et al., 2013; Tamnes et al., 2018). Notably, hippocampal subfields also display distinct levels of glucocorticoid receptors and show subfield-specific stress sensitivity (Christian et al., 2011; Gerlach and McEwen, 1972; McEwen, 1999; Wang et al., 2013).

Despite its complex structural organization, human neuroimaging studies have often focused on how youth adversity affects the overall hippocampal structure. Reductions in hippocampal volume were found in children exposed to socioeconomic disadvantage and childhood maltreatment (Botdorf et al., 2022; Hanson et al., 2015; Jednoróg et al., 2012; Luby et al., 2013, 2020; McLaughlin et al., 2019; Taylor et al., 2020; Teicher et al., 2018). However, treating the hippocampus as a uniform anatomical structure obscures distinct vulnerability patterns across hippocampal subregions. Indeed, subfield-specific susceptibility to adversity has been reported with greater reductions in CA4-DG and CA2-CA3 (Teicher et al., 2012). Along the hippocampal anterior-posterior axis, regional vulnerability is suggested to vary with the type and timing of adversity, involving different anterior and posterior regions (Teicher et al., 2018). Although evidence supports differential vulnerability across the anatomical organization of the hippocampus, it remains unclear whether these regional associations are consistently reflected in its complementary structural properties.

Addressing this question requires moving beyond volumetric measures. Other macrostructural features of the hippocampus may capture distinct aspects of organization and maturation in the context of adversity. Similar to the neocortex, the hippocampus exhibits variable gyrification patterns along its longitudinal axis, with considerable interindividual variability (DeKraker et al., 2018; Duvernoy, 1988; Morris et al., 2024). Other morphological features, such as thickness, also differ systematically across hippocampal subfields (DeKraker et al., 2020). Accounting for interindividual anatomical variability requires segmentation approaches that preserve hippocampal topology while aligning homologous tissue across individuals. HippUnfold is one such approach, which creates an unfolded two-dimensional topological space that preserves subfield boundaries and facilitates surface-based analyses analogous to neocortical surface reconstruction (DeKraker et al., 2018, 2020, 2022, 2023).

However, macrostructural features provide only one perspective on hippocampal organization, whereas microstructural measures may offer further information by characterizing tissue properties and myeloarchitecture (Karat et al., 2023). Nonetheless, the impact of youth adversity on the hippocampal microstructure remains relatively understudied. Previous work with diffusion MRI reported changes in microstructural properties of the hippocampus associated with childhood stress (Conley et al., 2021; McManus et al., 2024). Similarly, prematurity-related stress has been linked to decreased hippocampal myelin content inferred from T1w/T2w ratio and subfield-specific vulnerability (Nichols et al., 2025). However, prospective evidence on the association between ongoing youth adversity and hippocampal microstructure development remains limited.

This study investigated the associations between adversity in early adolescence and hippocampal macro- and microstructural features, namely volume, thickness, gyrification, and T1w/T2w ratio, an *in vivo* proxy for microstructure (Glasser and Van Essen, 2011). The hippocampus was analyzed as a whole structure and across subfields defined along its proximal–distal and anterior–posterior axes (Paquola et al., 2020). Adversity was assessed at the family and socioeconomic levels. Family-related adversity dimensions were derived from an Exploratory Factor Analysis (EFA) of caregiver- and youth-reported measures of family dynamics and parental style, whereas socioeconomic vulnerability was measured with the income-to-needs ratio and Area Deprivation Index, an indicator of neighborhood socioeconomic disadvantage. In order to capture both the burden of co-occurring adversity types and their distinct individual associations, we examined cumulative adversity using univariate models and individual adversity types using multivariate analyses that account for their shared covariance. We leveraged a wide population-based cohort of 9-11-year-olds (*N* = 5, 263) from the baseline assessment of the Adolescent Brain Cognitive Development (ABCD^®^) Study and used HippUnfold, an advanced automatic segmentation approach that derives surface-based hippocampal metrics while preserving interindividual differences (DeKraker et al., 2022). We hypothesize that hippocampal macro- and microstructural features in early adolescence differ as a function of exposure to adversity with subfield-specific signatures.

## Results

Adversity was characterized using both family-related and socioeconomic measures. Family-related adversity was summarized using Exploratory Factor Analysis (EFA) of caregiver- and youth-reported questionnaire items assessing family environment and parental behavior. The EFA identified three family-related adversity dimensions capturing youth-reported family environment, caregiver-reported family environment, and neglectful parenting. Socioeconomic measures were assessed through the Area Deprivation Index (ADI) and income-to-needs ratio (**Fig. 1A**). ADI reflects the socioeconomic status of a neighborhood, with higher values indicating greater deprivation. The income-to-needs ratio represents household income relative to the poverty threshold adjusted for household size. Sex, age, puberty, intracranial volume, and body mass index (BMI) were used as covariates in all models to account for their potential influences on development and hippocampal morphology (Brzezinski-Rittner et al., 2026; Hashimoto et al., 2015; Mestre et al., 2020; Mills et al., 2016; Satterthwaite et al., 2014).

**Figure 1.**
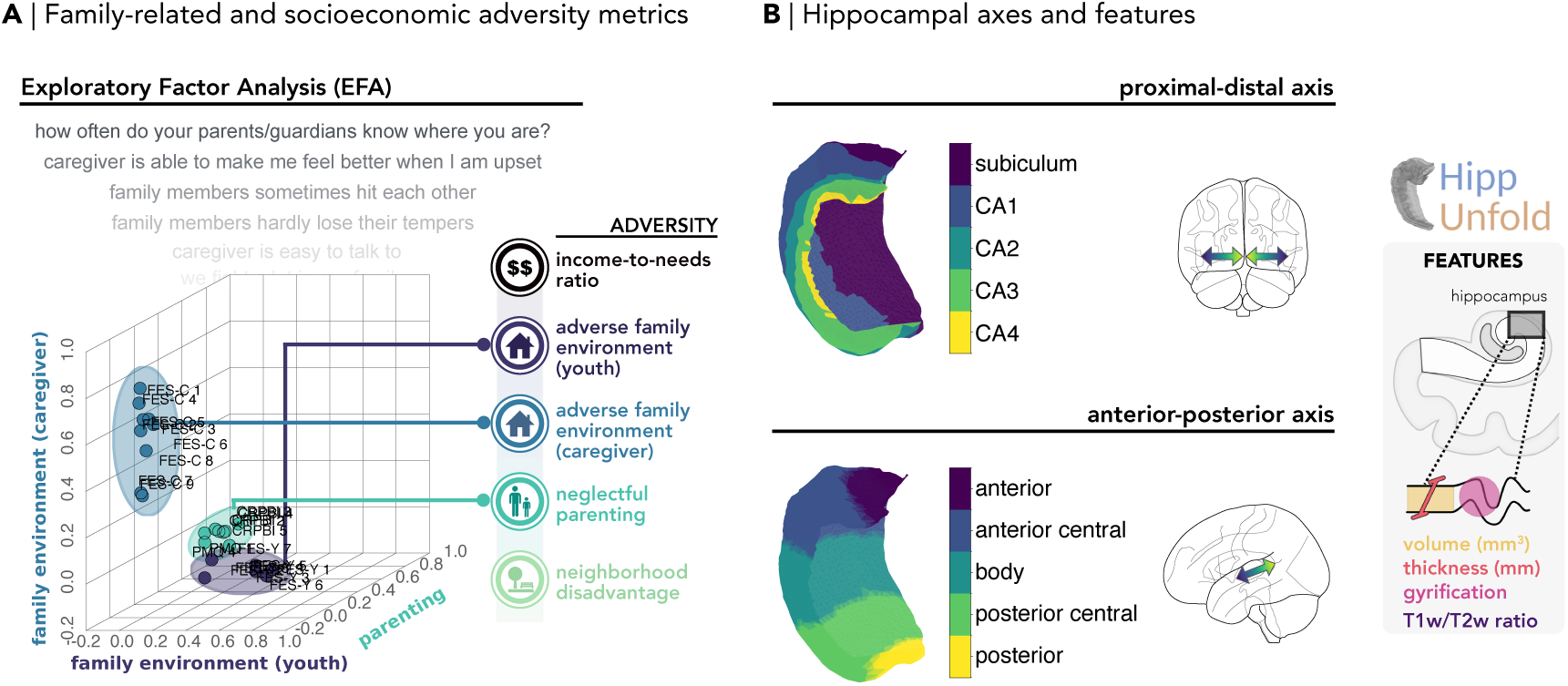
Overview of adversity metrics considered and hippocampal subfields investigated. **A** | Exploratory Factor Analysis (EFA) yielded a three-factor solution reflecting parenting and family environment reported by youth and caregiver. Socioeconomic measures were assessed using the income-to-needs ratio and neighborhood disadvantage measured by the Area Deprivation Index. **B** | Hippocampal subfields along the proximal-distal and anterior-posterior axes were analyzed at the macro- (volume, thickness, and gyrification) and microstructural (T1w/T2w ratio) levels.

HippUnfold was used to derive hippocampal macro- and microstructural features. Analyses were conducted at the whole-hippocampus and across the anterior-posterior and proximal-distal axes using volumetric and surface-based features (**Fig. 1B**). Volumetric analyses included the subiculum, cornu ammonis (CA) 1-4, stratum radiatum lacunosum moleculare (SRLM), and dentate gyrus (DG). Surface-based analyses included thickness, gyrification, and T1w/T2w ratio measures assessed across the proximal-distal subfields (subiculum and CA1-4) and five segments of the anterior-posterior axis (anterior, anterior central, body, posterior central, and posterior).

The final sample comprised 5,263 participants (mean age = 119 ± 7.55 months, 48% females; **Table 1**). To evaluate potential selection bias, factor scores derived from the EFA in the full questionnaire sample were compared between participants with and without usable MRI data (*N* = 8, 666), as well as those present in the final sample (*N* = 5, 263). Differences across all factors were negligible (all Cohen’s |*d*| < 0.05), indicating minimal impact of sample selection procedures on the latent constructs. The three-factor structure identified in the EFA in the full questionnaire sample was supported in the final sample, as indicated by a Confirmatory Factor Analysis (CFA) with overall good fit (CFI = 0.93, TLI = 0.93, RMSEA = 0.030, 90% CI [0.028, 0.032]).

**Table 1.** Description of sample characteristics in relation to its biological and developmental aspects.

| **Biological characteristics** | **Mean (SD)** |
| --- | --- |
| Age (months) | 119 (7.55) |
| BMI | 18.59 (3.95) |
| ICV (mm<sup>3</sup>) | 1,503,897 (142,404) |
| **Sex distribution** | **N (%)** |
| Male | 2,731 (51.9%) |
| Female | 2,532 (48.1%) |
| **Developmental characteristics** |  |
| Pubertal stage |  |
| 1 | 2,769 (52.6%) |
| 2 | 1,252 (23.8%) |
| 3 | 1,176 (22.3%) |
| 4 | 62 (1.2%) |
| 5 | 4 (0.1%) |

### Subfield- and axis-specific effects of cumulative adversity exposure

To investigate the effects of aggregated adversity types, cumulative adversity exposure was indexed as the sum of family-related adversity factor scores, neighborhood disadvantage, and reversed income-to-needs ratio, with higher values indicating greater adversity. Linear mixed-effects models assessed associations of cumulative adversity with hippocampal macro- and microstructural features at the whole-hippocampus and across subfields of both axes, while accounting for family structure and biological variables relevant to adolescent neurodevelopment (puberty, sex, intracranial volume, and BMI).

Overall, increased cumulative risk was associated with decreased hippocampal macrostructure at the whole-hippocampus and subfield-specific levels (non-standardized *β*_range_ = −13.59 to −0.001 across measures, all *p*_FDR_ < 0.05). Across volume, thickness, and gyrification, the subiculum and CA1 showed consistent negative association with cumulative adversity exposure (**Fig. 2A**) as well as with the whole hippocampus. Additionally, reduced volumes in the SRLM, CA2, CA3, and DG were associated with accumulated adversity. In the anterior-posterior axis, cumulative adversity was associated with decreases in thickness and gyrification at the extreme anterior and posterior ends. These effects extended to the anterior-central and posterior-central regions of the axis for gyrification and thickness, respectively. In contrast, positive associations with T1w/T2w ratio were restricted to the subiculum and posterior end of the hippocampus (non-standardized *β*_range_ = 0.004 to 0.005, all *p*_FDR_ < 0.05). Marginal and conditional *R*^2^ values for all linear mixed-effects models are reported in **Supplementary Table S1**. Together, these findings indicate that cumulative adversity was consistently associated with reductions in hippocampal volume, thickness, and gyrification, particularly within the subiculum and CA1, whereas significant positive associations were observed in the subiculum and posterior end for T1w/T2w ratio.

### Widespread spatial associations of hippocampal macrostructure with socioeconomic measures

We next investigated whether individual adversity types showed distinct associations with hippocampal macrostructure. To address this question, Partial Least Squares Regression (PLS-R) analyses were conducted separately for each macrostructural feature at the whole-hippocampus level and along the proximal-distal and anterior-posterior axes. Sex, puberty, intracranial volume, and BMI were considered alongside adversity predictors. Permutation testing and bootstrap resampling indicated that models with higher-order components were not stable and consistently recovered across samples for some hippocampal structural features. Thus, to ensure consistency and comparability across spatial scales, only the first component was retained for inference.

**Figure 2.**
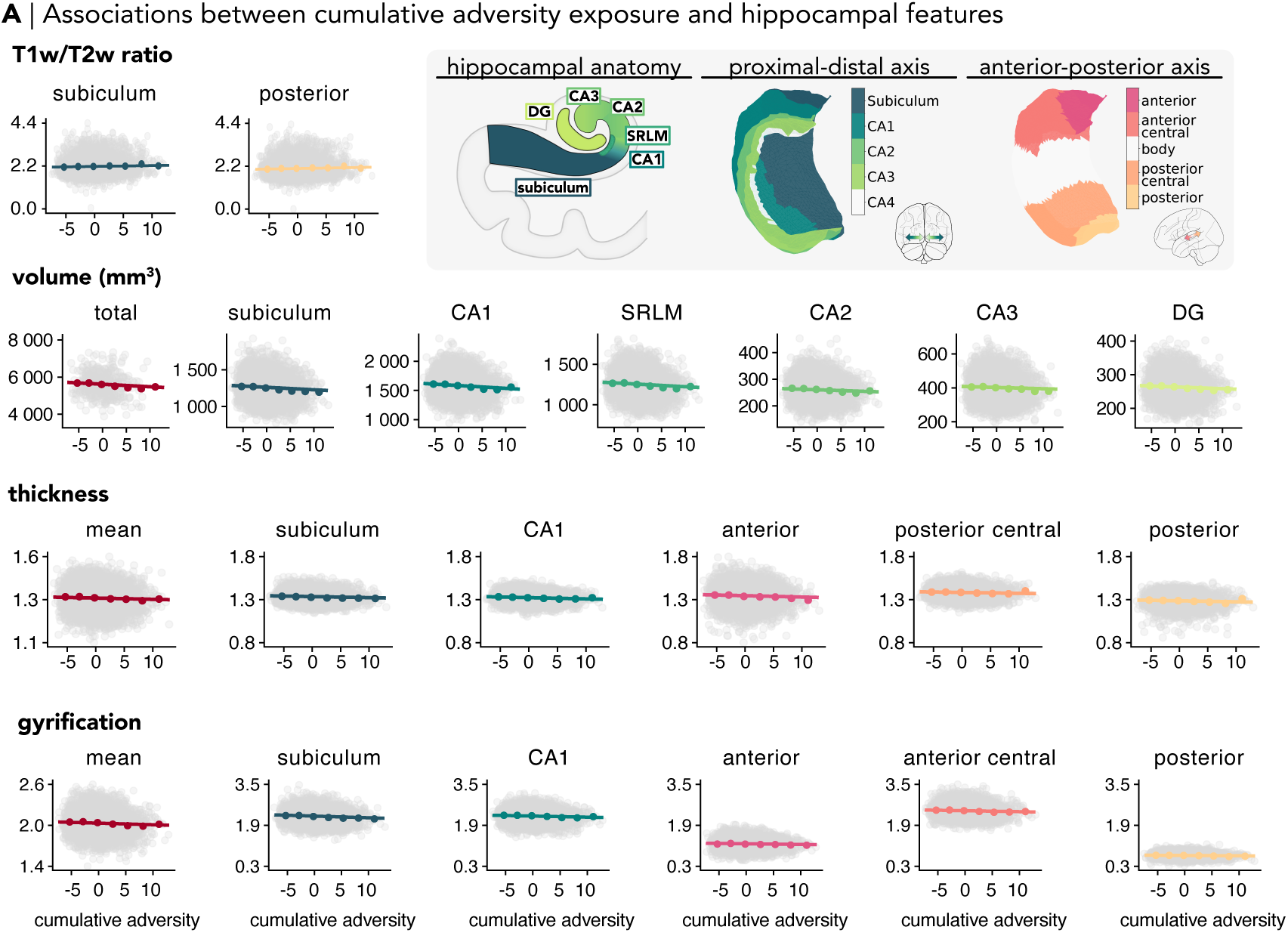
Cumulative adversity exposure associations across different hippocampal features. **A** | Cumulative adversity exposure (z-standardized) was associated with reductions of distinct subfields along the anterior-posterior and proximal-distal axes across all macrostructural features (volume, thickness, and gyrification). In contrast, cumulative adversity exposure was associated with increases in T1w/T2w ratio of the subiculum and posterior end of the hippocampus. Only results surviving FDR correction (*p*_FDR_ < 0.05) are shown. Lines represent fixed-effect predictions from linear mixed effects models, with variations of cumulative adversity while holding other covariates constant (continuous variables at 0, corresponding to the sample mean, and categorical variables at reference levels, e.g., males, puberty stage 1). Gray points represent individual observations. Filled circles represent the mean observed outcome within seven equally spaced bins of the standardized cumulative adversity exposure and are shown for visualization purposes only. Hippocampal schematics on the top right illustrate the spatial organization of the hippocampal subfields along the proximal-distal and anterior-posterior axes. Colored regions indicate statistically significant associations (*p*_FDR_ < 0.05), whereas white regions represent regions that were not statistically significant.

Across macrostructural models, the first component described a general sociodemographic-developmental dimension at both whole-hippocampus and axis levels, explaining between 10.63% and 35.70% of the variance across models (*p*_permutation_ < 0.001). Socioeconomic measures showed stronger loadings on the dominant component and higher bootstrapped VIP scores compared to family-related adversity indicators (**Fig. 3A**). Biological variables exhibited the highest VIP values; specifically, the lower bound of the bootstrapped confidence interval exceeded the conventional threshold for intracranial volume (ICV) and sex (95% CI of VIP > 1). Taken together, these results suggest that the latent component was jointly characterized by biological and socioeconomic predictors.

**Figure 3.**
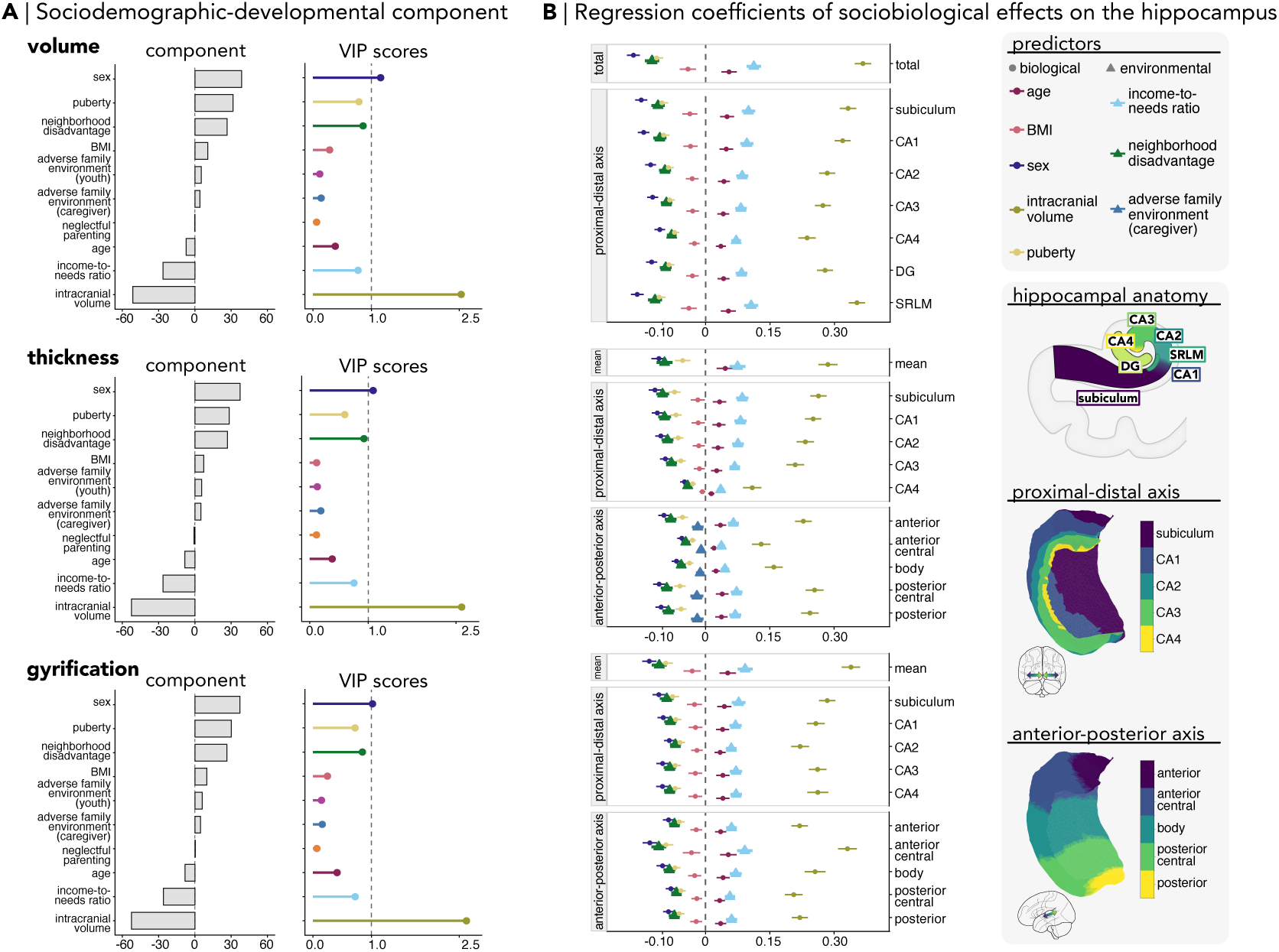
Associations between different adversity types and hippocampal macrostructure. **A** | First component of Partial Least Squares Regression (PLS-R) captured a sociodemographic-developmental dimension. Component loadings represent the contribution of each predictor to the first latent component. Variance Importance in Projection (VIP) scores indicate the relevance of each variable for predicting hippocampal measures. **B** | Regression coefficients of PLS-R showed associations of socioeconomic measures at the whole hippocampus and axes levels across all macrostructural features. Only results surviving FDR correction (*p*_FDR_ < 0.05) are shown. Points indicate PLS-R coefficients and horizontal error bars indicate 95% confidence intervals estimated from family-level resampling. Hippocampal schematics on the right illustrate the spatial organization of the different hippocampal subfields along the proximal-distal and anterior-posterior axes.

Socioeconomic measures were associated with volume, thickness, and gyrification at both whole-hippocampus and axis levels. Higher neighborhood disadvantage was associated with lower values across these macrostructural features (*β*_range_ = −0.12 to −0.04, all *p*_FDR_ < 0.001), whereas higher income-to-needs ratio predicted greater volume, thickness, and gyrification (*β*_range_ = 0.04 to 0.11, all *p*_FDR_ < 0.001; **Fig. 3B**). In contrast, family-related adversity showed an axis-specific effect restricted to hippocampal thickness. More specifically, adverse family environment reported by caregivers was associated with decreased thickness of the subregions along the anterior-posterior axis (*β*_range_ = −0.02 to −0.01, all *p*_FDR_ = 0.012 − 0.015). Additional model performance metrics are provided in **Supplementary Table S2**. Overall, socioeconomic measures exhibited widespread associations across hippocampal macrostructure. In contrast, family-related adversity showed associations restricted to hippocampal thickness along the anterior-posterior axis.

### Spatially distinct associations of hippocampal T1w/T2w ratio with neighborhood disadvantage

Next, we examined whether the macrostructural patterns extended to hippocampal microstructure. PLS-R models of T1w/T2w ratio were fitted using the same predictors as in the macrostructural analyses, i.e., the effects of family-related and socioeconomic measures were assessed while accounting for biological features. Consistent with the macrostructural analyses, only the first component was retained for inference to ensure stable model estimation across spatial scales.

The first component of the PLS-R of T1w/T2w ratio described a general sociodemographic-developmental dimension at the whole hippocampus and axis levels (component explained variance range = 2.39-2.50%, *p*_permutation_ < 0.001; **Fig. 4A-i**). Although socioeconomic variables contributed to the component, biological variables, particularly BMI and intracranial volume, showed the highest importance (95% CI of VIP > 1), indicating that the latent component captured variation related to both biological and socioeconomic predictors. Spatially, significant associations with socioeconomic measures were observed only along the anterior–posterior axis, where neighborhood disadvantage was associated with increased T1w/T2w ratios across all subregions (*β* = 0.03, all *p*_FDR_ = 0.04; **Fig. 4A-ii**). In contrast to findings in macrostructure, microstructural associations were limited to neighborhood disadvantage along the anterior-posterior axis.

**Figure 4.**
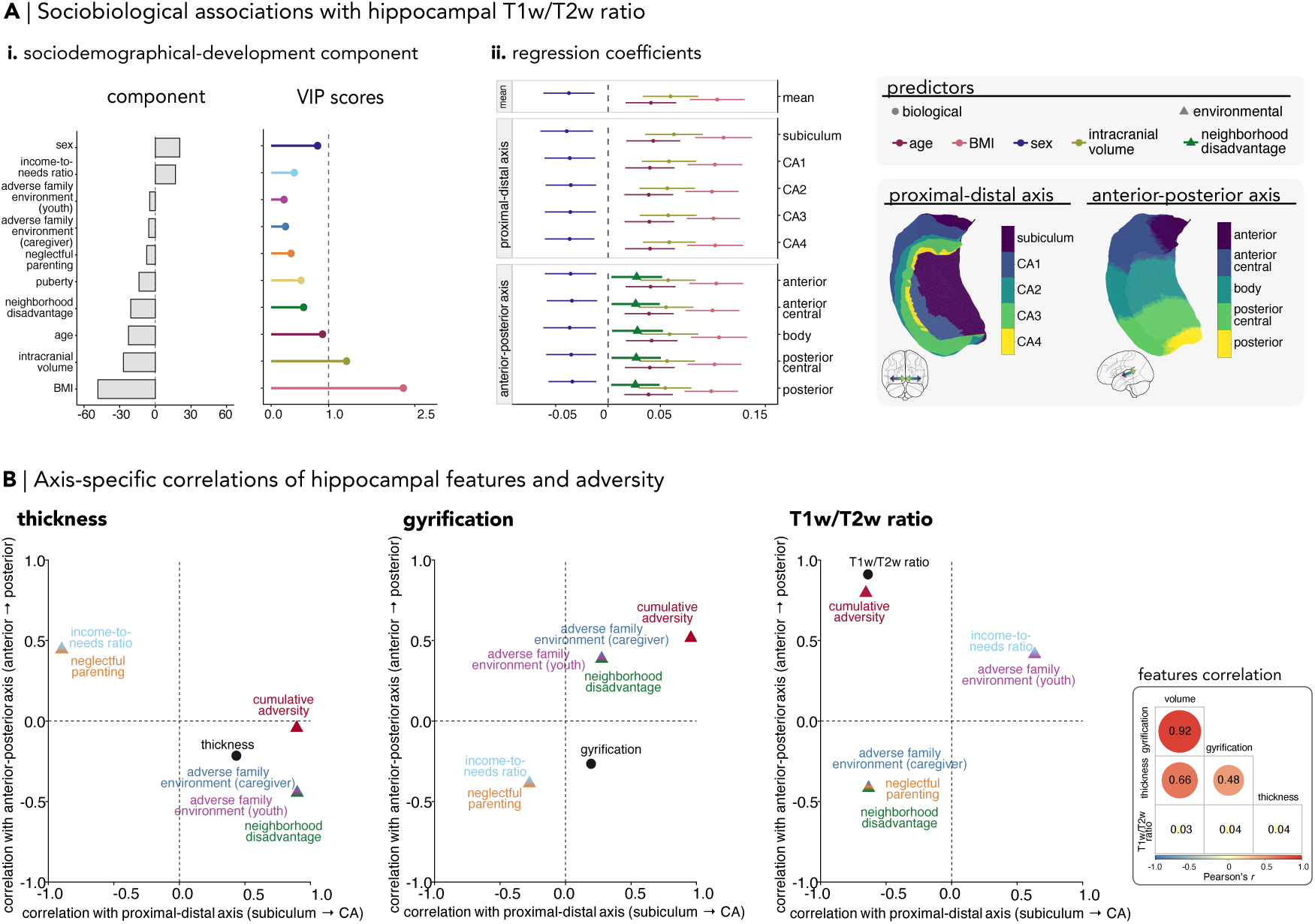
Associations of adversity with hippocampal T1w/T2w ratio. **A** | **i**. The first component of Partial Least Squares Regression (PLS-R) captured a sociodemographic-developmental dimension. Component loadings represent the contribution of each predictor to the first latent component. Variance Importance in Projection (VIP) scores indicate the relevance of each variable for predicting hippocampal measures. **ii**. Regression coefficients of PLS-R showed effects of neighborhood disadvantage along the hippocampal anterior-posterior axis. Only results surviving FDR correction (*p*_FDR_ < 0.05) are shown. Points indicate PLS-R coefficients and horizontal error bars indicate 95% confidence intervals estimated from family-level resampling. **B** | Axis-specific Pearson correlations descriptively summarize the spatial organization of hippocampal features and adversity effects. Correlations were computed between subfield position along the proximal–distal and anterior–posterior axes and either mean feature values or regional effect estimates. The effect estimates of all adversity types were used irrespective of their statistical significance. Each point represents the strength and direction of association along the two axes. Positive values indicate increasing values toward posterior or distal (CA) regions, whereas negative values indicate increasing values toward anterior or proximal (subiculum) regions. On the right, subject-level correlations between summed hippocampal volume and average hippocampal thickness, gyrification, and T1w/T2w ratio.

To visualize the spatial organization of hippocampal features and their associations with adversity along the proximal-distal and anterior-posterior axes, we computed descriptive Pearson correlations between subfield position (treated as equally spaced locations from 1 to 5) and group-mean feature values across the five regions of each axis. The same approach was applied to model-derived effect estimates of each adversity measure per subfield (**Fig. 4B**). These correlations were intended to summarize linear spatial trends for visualization and were not used for statistical inference. Among hippocampal features (represented as circles), T1w/T2w ratio showed stronger spatial gradients along both axes compared to thickness and gyrification, with values increasing toward the subiculum and posterior hippocampus. On the other hand, adversity effects (depicted as triangles) presented a heterogeneous, feature-specific spatial organization. In thickness and gyrification, the effect estimates of adversity showed a similar distribution along the proximal-distal axis, reflecting both subiculum- and CA-weighted patterns. Their anterior-posterior gradient was, however, opposite in direction across the two features, indicating that adversity may relate differently to hippocampal thickness and gyrification along the anterior-posterior axis. Subject-level correlations among hippocampal features showed high correspondence between volume and gyrification, moderate correspondence with thickness, and minimal correspondence with T1w/T2w ratio (**Fig. 4B**, right), illustrating the differing relationships among the hippocampal structural features.

## Discussion

In a large population-based cohort of 9-11-year-olds, adversity was associated with alterations in hippocampal macro- and microstructural features at the whole-structure and axis levels. The spatial distribution of the effects varied across adversity types, as revealed in both multivariate and univariate analyses. Socioeconomic measures showed widespread associations across hippocampal macrostructural features, while the effects on microstructure were restricted to the anterior-posterior axis. Associations between cumulative adversity exposure and hippocampal macro- and micro-structure were primarily characterized by subfield-specific patterns, with the subiculum and CA1 emerging as common affected regions in analyses of volume, thickness, and gyrification.

The latent component derived from the multivariate analyses reflects a developmental dimension intertwined with socioeconomic context. In contrast to family-related adversity, socioeconomic measures consistently contributed to this dimension across all hippocampal features, indicating that variation in macro- and microstructure may be systematically related to socioeconomic context during maturation. Notably, income-to-needs ratio and neighborhood disadvantage were the dominant adversity-related contributors to the latent component, which suggests that hippocampal structural organization may be influenced by socioeconomic measures at multiple ecological levels. Importantly, these environmental associations were observed alongside strong contributions from biological predictors, particularly intracranial volume, puberty, and sex, indicating that socioeconomic influences on hippocampal structure should be interpreted within this broader biological context. Such an interplay between maturation and socioeconomic context is in line with prior work linking altered neurodevelopmental trajectories to low socioeconomic status at both the household and neighborhood levels (Rakesh and Whittle, 2021; Rakesh et al., 2023). In fact, distinct hippocampal trajectories have been linked to socioeconomic status in longitudinal studies (Barch et al., 2020; Ellwood-Lowe et al., 2018; McDermott et al., 2019). The latent dimension found in the present study, thus, supports the use of multivariate approaches to capture environmental complexities, particularly during adolescence, when intricate social experiences span family, peer, and neighborhood contexts (Choudhury et al., 2023; Ferschmann et al., 2022).

Despite their shared covariance within the latent component, different adversity types showed distinct spatial associations with hippocampal macrostructure. Specifically, the income-to-needs ratio and neighborhood disadvantage showed extensive associations with hippocampal macrostructure at the whole structure and axis levels. Family environment, on the other hand, was spatially linked to the anterior-posterior axis. These differences may reflect the distinct ecological nature of the adversity types. Whereas family-related adversities may represent more proximal interpersonal stressors, socioeconomic measures can indicate limitations in access to multiple resources relevant for development (Bradley and Corwyn, 2002). For example, socioeconomically disadvantaged adolescents may be more exposed to food insecurity and poorer diet (McLaughlin et al., 2012; Pechey and Monsivais, 2016), unmet medical and mental health care (Newacheck et al., 2003), and unequal educational opportunities (Bradley and Corwyn, 2002; Evans, 2004). The diffuse spatial associations observed for socioeconomic measures may, thereby, reflect the risk multiplicity for hippocampal maturation of distinct environmental domains embedded in socioeconomic conditions.

Our findings further suggest that the hippocampal macro- and microstructural features capture relatively distinct aspects of hippocampal maturation. Not only environmental factors, but also biological predictors contributed differently to the effects observed on macro- and microstructure. Income-to-needs ratio and puberty, for example, were related to hippocampal volume, thickness, and gyrification, while these associations were not significant in the T1w/T2w ratio. Previous work has similarly reported dissociable patterns between macro- and microstructure. While socioeconomic and family-related adversity has often been implicated in alterations of cortical and subcortical macrostructure (Alnæs et al., 2020; Breslin et al., 2024; Hanson et al., 2015; Rakesh and Whittle, 2021; Rao et al., 2010; Thorsen et al., 2025; Whittle et al., 2013), its association with T1w/T2w ratio has not been consistently observed (Boroshok et al., 2023; Langensee et al., 2022; Norbom et al., 2022; Thorsen et al., 2025; Weissman et al., 2023). In the hippocampus, underlying cellular and synaptic processes have been linked to macrostructural alterations to some extent (Czéh and Lucassen, 2007; Schoenfeld et al., 2017). However, previous work indicates that hippocampal macro- and microstructure offer complementary rather than redundant information on aging trajectories (Langnes et al., 2020; Ma et al., 2026; Pereira et al., 2014; Wolf et al., 2015). Here, we expand these findings to the effects of environmental context on different hippocampal features, suggesting that socioeconomic conditions may relate to the hippocampal macro- and microstructure through different biological processes.

Because youth adversity exposures often co-occur, cumulative scores summing up different adversity types have therefore been used. Such approaches are useful for identifying children exposed to high levels of environmental risk, which have been associated with concurrent and prospective effects on physical and mental health as well as cognitive development (Appleyard et al., 2005; Arata et al., 2007; Felitti et al., 1998; Evans et al., 2013). Subfield-specific associations were observed in the present study when adversity was operationalized as a cumulative index. Most notably, alterations in the subiculum and CA1 were consistent across macrostructural features, indicating potential sensitivity of those regions to cumulative stress across multiple aspects of hippocampal morphology. Rodent models further support associations between stress-related processes and CA1. For instance, prolonged administration of glucocorticoids led to alterations in the pyramidal cells of CA1 (Kadar et al., 1998). This subfield has a high density of glucocorticoid receptors, which can modulate glutamatergic transmission in the hippocampus (Joëls, 2009). Notably, CA1 neurons exhibit an elevated number of NMDA receptors, whose sustained activation promotes a high influx of calcium ions and may contribute to the sensitivity of CA1 to stress-related alterations (Stanika et al., 2010). Given that CA1 and subiculum compose the primary output regions of the hippocampus and are central components of the regulation of the HPA-axis via projections to limbic and hypothalamic structures (Herman et al., 2016; MacDougall and Howland, 2013), morphological alterations in these regions may therefore reflect stress-related modulation of hippocampal regulatory pathways associated with cumulative adversity exposure.

The observed associations should take into consideration the ongoing developmental context of the hippocampus. The hippocampal proximal-distal and anterior-posterior axes show distinct maturational patterns, with region-specific age-related structural changes (Daugherty et al., 2016, 2017; DeMaster et al., 2014; Gogtay et al., 2006; Langnes et al., 2020; Lin et al., 2013; Mareckova et al., 2025; Riggins et al., 2018; Tamnes et al., 2018). In particular, the anterior region of the anterior-posterior axis exhibits protracted maturational trajectories (Langnes et al., 2020; Riggins et al., 2018) and co-develops with large-scale cortical systems involved in higher-order cognition (Plachti et al., 2023). These age-related differences underscore the importance of interpreting the present findings within the developmental context of the current sample. The relatively narrow age range reduced developmental heterogeneity and provided a more focused characterization of environmental and biological associations during early adolescence.

Although the findings of the present study shed light on the association between adversity and hippocampal macro- and microstructure during early adolescence, the directionality of this relationship remains an open question. Most interpretations in the literature assume that adversity may shape hippocampal morphology (McLaughlin et al., 2019; Rakesh and Whittle, 2021). It may also be plausible that pre-existing differences in hippocampal structure interact dynamically with environmental conditions, influencing individual sensitivity to adversity. Animal models, specifically, have identified several molecular and functional hippocampal markers associated with neurobiological adaptation to stress (Larosa and Wong, 2022). In humans, youth considered resilient following adversity exposure have also been linked to differences in hippocampal volume (Morey et al., 2016), as well as to functional and neuromodulatory processes involving prefrontal-hippocampal circuitry (Wei et al., 2025). It is thereby relevant to note that adversity-related alterations in the hippocampus may emerge through dynamic and potentially adaptive mechanisms, rather than solely reflecting a simple unidirectional pathway of vulnerability. Disentangling the directionality of the adversity-hippocampus relationship will therefore remain an important challenge for future developmental research.

It is also relevant to consider the methodological limitations of the current study. First, some demographic variables were recommended careful handling, particularly household income, due to systematic patterns of missingness reported in the ABCD documentation. Declined responses and “Do Not Know” answers were not randomly distributed across participants and may correlate with other sociodemographic variables. Within the sample of participants with usable MRI data, most “Do Not Know” responses were reported in Hispanic households, whereas “Decline to Answer” responses were more frequently observed among White participants, consistent with patterns described in the ABCD documentation (**Supplementary Table S3**). After applying our exclusion criteria, the ethnic distribution of the final sample remained broadly comparable to that of the MRI-eligible sample and to patterns reported in the ABCD documentation, suggesting that our selection procedures did not substantially introduce additional selection bias beyond that present in the dataset (**Supplementary Table S4**). It is worth noting that the construction of the Area Deprivation Index (ADI) derived from the Neighborhood Atlas has been questioned. Specifically, Hannan et al. (2023) reported that home value- and income-related variables may be disproportionately represented in ADI scores, requiring, therefore, careful consideration of the composite index. In the context of hippocampal feature extraction, HippUnfold may underestimate the complexity of hippocampal gyrification, particularly at 1 mm isotropic resolution. Accordingly, follow-up studies using high-resolution, hippocampal-targeted structural imaging may provide more sensitive estimates of this feature. Other methodological limitations concern the interpretation of T1w/T2w ratio measures. Although T1w/T2w ratio has been widely used as an *in vivo* proxy for myeloarchitectural organization (Chao et al., 2015; Karat et al., 2023; Langensee et al., 2022; Nichols et al., 2025; Thorsen et al., 2025; Weissman et al., 2023), its biological meaning remains complex. Previous work comparing T1w/T2w ratio to histologically validated myelin measures has generally reported low-to-moderate correlations, which suggests that it reflects multiple microstructural tissue properties beyond myelin content (Arshad et al., 2017; Uddin et al., 2018, 2019). In particular, T1w/T2w ratio can be influenced by additional tissue properties, such as iron (Stüber et al., 2014) and water concentration (Miot-Noirault et al., 1997). Furthermore, bias-field correction may be warranted to reduce intensity inhomogeneities that could obscure T1w/T2w ratio distributions, particularly in multisite datasets (Nerland et al., 2021). However, preprocessing approaches intended to improve spatial correspondence and inter-subject comparability may also influence biologically meaningful interindividual variation in T1w/T2w estimates. Lastly, the absence of radiofrequency transmit field (B1 +) corrections for the T1w/T2w ratio, as proposed by Glasser et al. (2022), may have allowed residual transmit field biases to influence the microstructural estimates in the current study. Therefore, methodological choices related to T1w/T2w preprocessing should be carefully considered when interpreting individual and group-level differences.

The current study demonstrates that different types of adversity are associated with distinct hippocampal structural properties and spatial patterns, highlighting the multidimensional relationship between environmental context and hippocampal maturation. By jointly investigating macro- and microstructural features and the two principal hippocampal axes in a large population-based cohort of early adolescents, the present study provides a comprehensive spatial framework for understanding how different adversity types are linked to hippocampal organization. Socioeconomic conditions, family-related adversity, and cumulative adversity exposure relate selectively to macrostructural and microstructural features, suggesting complementary patterns of hippocampal organization rather than a single adversity-related phenotype. Such observations, therefore, highlight the importance of conceptualizing both the environment and the hippocampal structure as multidimensional systems during development.

## Methods

### Sample

Data were obtained from the Adolescent Brain Cognitive Development (ABCD^®^) Study (5.1 release). Analyses were based on ABCD baseline data, including behavioral, demographic, and neuroimaging measures. Only one participant from multiple-birth families (twins or triplets) was considered to ensure independence of observations; non-twin siblings were nonetheless included.

### Behavioral and anthropometric data

#### Assessment of family-related adversity

Questionnaires at baseline on the youth’s social environment were chosen to quantify adversity. Assessment of family social climate was obtained with the Family Environment Scale of the PhenX Toolkit (FES; Hendershot et al., 2015; edition of 1994, as reported in the ABCD Data Documentation) - Family Conflict subscale, due to a higher number of respondents. This subscale evaluates self-reported perception of open hostility and aggression within a family (Rivers and Sanford, 2018). Responses of the youth and caregiver were separately included in the analysis due to low agreement (Cohen’s kappa < 0.20) between caregivers’ and youth’s answers for all responders (*N* = 11, 832) and the final sample (*N* = 5, 263; **Supplementary Tables S5 and S6**). A similar low correspondence in items of the Family Environment Scale was previously reported within the ABCD study (Hogan et al., 2023).

Parental Monitoring Questionnaire (PMQ) and Children’s Report of Parental Behavior Inventory (CRPBI) were used to evaluate the youth-caregiver relationship. The former measures communication between caregivers and the youth, as well as caregivers’ awareness of the youth’s whereabouts (Karoly et al., 2016). The latter comprises the Acceptance Scale of the CRPBI, which estimates emotional support and caregiver warmth (Schaefer, 1965).

As a complementary part of sample selection, participants were required to have non-missing responses to selected items of the Kiddie Schedule for Affective Disorders - Post-Traumatic Stress Disorder module (KSADS-PTSD) assessing exposure to severe adversities (e.g., physical or sexual violence; **Supplementary Table S7**). Although these variables were not analyzed due to their low prevalence, their inclusion as a selection criterion indicated that the majority of the sample did not report exposure to these events.

#### Assessment of socioeconomic measures

Income-to-needs ratio and the percentile of the Area Deprivation Index (ADI) were considered socioeconomic measures. ADI is a census tract composite score of 17 items measuring neighborhood socioeconomic status in which higher values represent greater disadvantage (Fan et al., 2021). The income-to-needs ratio was calculated by converting the ABCD combined household income to the midpoint of its bin and dividing this value by the poverty threshold provided by the U.S. Department of Health and Human Services (HHS) Poverty Guidelines corresponding to the year of the interview (2016-2018). Because the poverty threshold depends on the household size, participants with implausible household sizes were excluded.

#### Exploratory Factor Analysis (EFA)

To investigate family-related adversity, an exploratory factor analysis (EFA) was conducted with the questionnaire items of the FES (youth- and caregiver-reported), PMQ, and CRPBI with all respondents (*N* = 11, 505). Analyses were performed in R version 4.5.0 using the *psych* package (R Core Team, 2025; Revelle, 2025). Inspection of the inter-item correlation matrix indicated adequate correlations (r > 0.30) for factor analysis. Bartlett’s test of sphericity was significant, and the Kaiser-Meyer-Olkin (KMO) values ranged from 0.77 to 0.94, indicating ‘good to marvelous’ sampling adequacy (Field et al., 2012; Tabachnick and Fidell, 2013).

Due to the ordinal and binary nature of the questionnaire items, a Kendall-based correlation matrix was computed. Factors were extracted using weighted least squares estimation and oblique (promax) rotation, allowing correlation between factors and reflecting a more realistic representation of psychological constructs (Fabrigar et al., 1999; Goretzko et al., 2021). The number of retained factors was determined by scree plots, parallel analysis, eigenvalues > 1, and theoretical interpretability, converging to the retention of three factors (Field et al., 2012; Tabachnick and Fidell, 2013; **Supplementary Figure S1**).

A three-factor solution accounted for 36.2% of the total variance (Factor 1 = 13.1%, Factor 2 = 11.2%, Factor 3 = 11.9%; model fit index = 0.78) and yielded dimensions reflecting Parental Neglect, Youth-reported Family Environment, and Caregiver-reported Family Environment. To improve factor discrimination, items with loadings below 0.32 (∼10% of shared variance) were removed during model refinement (Tabachnick and Fidell, 2013), increasing the explained variance to 39.5% (Factor 1 = 13.2%, Factor 2 = 13.2%, Factor 3 = 13.2%; model fit index = 0.81). One retained item had a final loading of 0.302 after re-estimation of the final factor solution. Residual covariances were allowed between items with similar wording within the same questionnaire. Factor correlations and internal consistency (Cronbach’s alpha) are described in the **Supplementary Table S8**. This resulting factor structure was subsequently evaluated using Confirmatory Factor Analysis (CFA) in the final subsample to assess its fit within the participants included in the imaging analyses.

### Imaging procedure

#### MRI assessment

Structural magnetic resonance images (MRI) were acquired with similar sequences on 3T General Electric, Philips, or Siemens scanners. To reduce motion-related artifacts in pediatric imaging, participants were tested with a scanner simulation and trained for motion compliance. Motion correction was applied during the acquisition of T1- and T2-weighted images (T1w and T2w, 1mm isotropic) by the ABCD Data Analysis and Informatics Center (DAIC). Further information on scan protocol and MRI acquisition can be found in Casey et al. (2018) and Hagler Jr et al. (2019).

Baseline structural MRI data from the ABCD Fast-Track release were accessed through the Developmental Cognition and Neuroimaging (DCAN) Labs ABCD-BIDS Community Collection (ABCC). In addition to the quality control performed by the ABCD DAIC, participants with clinical findings or failing ABCD manual quality control for T1w and T2w images were excluded. Distortion correction and alignment of T1w and T2w images were performed using the DCAN ABCD-adapted version of the PreFreeSurfer step of the Human Connectome Project (HCP) processing pipeline (Glasser et al., 2013). Modifications to the HCP pipeline include nonlinear registration implemented in Advanced Normalization Tools (ANTs) and N4 Bias Field Correction (DOI 10.5281/zenodo.15200276).

#### Hippocampal segmentation

The hippocampus was segmented with the HippUnfold toolbox (v1.5.1), which performs an automated tissue segmentation using a U-Net deep convolutional neural network and reconstructs surface-based metrics through a topology-preserving unfolded coordinate space. This coordinate system enables inter-subject alignment despite individual morphologies and exhibits improved subfield segmentation compared to conventional registration methods (DeKraker et al., 2022, 2023).

The unfolded space was used to segment subfields in the proximal-distal (PD) axis based on the multihist7 atlas (DeKraker et al., 2023), resulting in the division of the hippocampus into the subiculum, CA1, CA2, CA3, CA4, stratum radiatum lacunosum (SRLM), and dentate gyrus (DG). In this configuration, the subiculum is the most proximal region to the neocortex. We then divided the hippocampal subregions along the anterior-posterior (AP) axis based on the multihist7 atlas to apply the DeKraker25 parcellation, with 5 x 5 parcels spanning the PD and AP axes. Subregions in the AP axis were defined as anterior, anterior central, body, posterior central, and posterior by averaging five parcels along the PD axis to the corresponding anterior-posterior parcel. Participants with segmentation DICE overlap < 0.70 were excluded (DeKraker et al., 2022).

#### Hippocampal features

We assessed the effects of adversity on three macrostructural features (volume, thickness, and gyrification) and T1w/T2w ratio, as a proxy for microstructure (**Supplementary Figure S2**). The T1w/T2w ratio maps were generated with HippUnfold by dividing bias-corrected T1-weighted images by the corresponding T2-weighted images. Due to differences in T1 and T2 relaxation times, lipid-rich content appears brighter in T1w and darker in T2w images. This contrast difference is enhanced by the T1w/T2w ratio, which can serve as a proxy for myelin-sensitive content (Glasser and Van Essen, 2011).

Thickness, gyrification, and T1w/T2w ratio measures were projected onto the surface and averaged along the regions of the AP and PD axes. Surface estimates of the DG have low reliability at a 1 mm isotropic resolution due to the small size and high folding of this subfield. Therefore, the DG was not included in the surface-based analysis of thickness, gyrification, and T1w/T2w ratio. Whole hippocampal measures were derived by aggregating included subfield values. Whole hippocampal volume corresponded to the sum of all subfield volumes (subiculum, CA1-4, SRLM, DG), whereas thickness, gyrification, and T1w/T2w ratio corresponded to the mean across hippocampal subfields (subiculum, CA1-4).

#### Harmonization

MRI scanning across different sites may introduce non-biological biases in the data, which can be mitigated through harmonization (Fortin et al., 2018; Saragosa-Harris et al., 2022). ComBat is a harmonization method developed to remove unwanted sources of variability in the mean and variance while preserving covariate effects (Fortin et al., 2018; Johnson et al., 2007).

To reduce potential bias in the estimation of harmonization parameters, extreme outliers were excluded prior to harmonization. Outliers were defined as values exceeding ± 4 standard deviations from the mean for each scanner, resulting in the exclusion of nine participants scanned on a General Electric system, seven on a Philips system, and 36 on a Siemens system.

To prevent data leakage in the harmonization step and subsequent analyses, we first divided the sample into 70% training and 30% test sets, while assigning family members to the same subset. NeuroHarmonize is an extension of ComBat that enables model parameters to be estimated from the training data and subsequently applied to the test set, thereby ensuring that site effects are corrected to the same reference distribution (Pomponio et al., 2020). This harmonization step was applied independently across the AP and PD axes and implemented in Python v3.13.1.

Despite harmonization, T1w/T2w ratio values from the Philips scanner remained systematically higher than those obtained from the General Electric and Siemens scanners (**Supplementary Figure S3**). To avoid residual scanner-specific bias, participants scanned with the Philips system were excluded from subsequent analyses. Re-estimation of harmonization and all downstream analyses were therefore conducted using data acquired on General Electric and Siemens scanners only. Importantly, the exclusion of these participants did not significantly alter all the sample characteristics (**Supplementary Figure S3**).

#### Covariates

Intracranial volume (ICV), age, sex assigned at birth, caregiver-reported pubertal stage from the Pubertal Development Scale (PDS), and body mass index (BMI) were included as covariates in all analyses to account for developmental, anatomical, and physiological sources of variability (Brzezinski-Rittner et al., 2026; Dong et al., 2021; Hashimoto et al., 2015; Kilpatrick et al., 2023; Mestre et al., 2020; Mills et al., 2016; Satterthwaite et al., 2014). ICV estimates were obtained from the tabulated ABCD Study variables derived through the standardized FreeSurfer-based structural MRI processing pipeline described previously (Hagler Jr et al., 2019). Age corresponded to the participant’s age at interview and was expressed in months. Puberty was assessed using the Pubertal Development Scale (PDS), a four-point Likert questionnaire assessing perceived physical maturation through common and sex-specific items (Petersen et al., 1988). Pubertal category scores were derived from summed PDS item scores, resulting in classifications of prepubertal, early pubertal, midpubertal, late pubertal, and postpubertal stages. Further details regarding PDS scoring procedures are described elsewhere (Cheng et al., 2021). Caregiver-reported PDS scores were used due to their higher reliability during childhood and adolescence (Beltz et al., 2026). BMI was calculated from measured height and weight. Biologically implausible anthropometric values were identified using the CDC growth chart algorithm implemented in *cdcanthro*, following the 2016 CDC modified z-score thresholds for height-for-age (*mod haz < −*5 or > 4), weight-for-age (*mod waz < −*5 or > 8), and BMI-for-age (*mod bmiz < −*4 or > 8). Participants were retained only if all three values were within the plausible range (Freedman et al., 2015).

### Modeling

All modeling analyses were conducted in the final sample, defined as participants with quality-controlled structural MRI data and complete adversity and covariate information. The final sample was also used for a Confirmatory Factor Analysis (CFA).

#### Effects of individual adversity

Partial Least Squares Regression (PLS-R) was applied to investigate the effects of individual adversity types while accounting for their shared covariance. In this framework, PLS-R identifies latent components that maximize the covariance between predictor and outcome matrices (Krishnan et al., 2011; Durham et al., 2023). Predictor and outcome matrices were standardized prior to model estimation, as implemented in the R package *mdatools* (Kucheryavskiy, 2020). Separate PLS-R models were estimated for whole-hippocampus, proximal-distal axis, and anterior-posterior axis analyses. Whole-hippocampus models included a single outcome variable, whereas axis-specific models included multiple regional measures corresponding to hippocampal subfields (proximal-distal axis) or anterior-posterior regions (anterior-posterior axis).

Model complexity was initially assessed using 10-fold family-level cross-validation within the training data. Given instability of higher-order components in some models, subsequent inference focused on the first latent component. Statistical significance of latent components was then evaluated with family-block permutation testing (b = 1000). For valid inferences in the presence of family structure, regression coefficients and Variable Importance in Projection (VIP) scores were estimated using family-level bootstrapping (b = 1000).

Model coefficients were divided by the standard error of their bootstrap distributions to compute z-statistics, and two-sided p-values were derived from the standard normal distribution. P-values were subsequently corrected for multiple comparisons using a false discovery rate (FDR) across all coefficient tests. Normal and basic bootstrap confidence intervals were also computed for VIP scores, with variables classified as important when the lower confidence bound exceeded 1 (Afanador et al., 2014).

#### Cumulative adversity exposure

Measures of adversity were z-standardized and summed to compose the cumulative adversity exposure score, computed with family-adversity factor scores, neighborhood disadvantage, and reversed income-to-needs ratio, with higher values indicating greater adversity. In contrast to PLS-R models, hippocampal outcomes were analyzed in their original harmonized scale. Continuous covariates were z-standardized prior to model estimation, whereas categorical variables (sex and pubertal stage) were included as factors. Linear mixed models were fit separately for each hippocampal region (AP and PD subfields and global measures), while taking into account covariates and family structure, as follows:

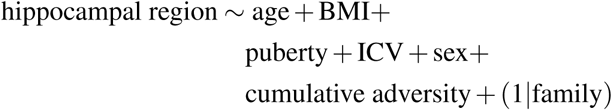

P-values were adjusted according to the global FDR correction across all fixed-effect tests pooled across all mixed models.

## Data availability

Data used in the preparation of this article were obtained from the Adolescent Brain Cognitive Development^SM^ (ABCD) Study, held in the NIMH Data Archive (NDA). This is a multisite, longitudinal study designed to recruit more than 10,000 children age 9-10 and follow them over 10 years into early adulthood. The ABCD Study^®^ is supported by the National Institutes of Health and additional federal partners under award numbers U01DA041048, U01DA050989, U01DA051016, U01DA041022, U01DA051018, U01DA051037, U01DA050987, U01DA041174, U01DA041106, U01DA041117, U01DA041028, U01DA041134, U01DA050988, U01DA051039, U01DA041156, U01DA041025, U01DA041120, U01DA051038, U01DA041148, U01DA041093, U01DA041089, U24DA041123, U24DA041147. A full list of supporters is available at Federal Partners - ABCD^®^ Study. A listing of participating sites and a complete listing of the study investigators can be found at Consortium Members - ABCD^®^ Study. ABCD consortium investigators designed and implemented the study and/or provided data, but did not necessarily participate in the analysis or writing of this report. This manuscript reflects the views of the authors and may not reflect the opinions or views of the NIH or ABCD consortium investigators.

The ABCD data repository grows and changes over time. The ABCD behavioral data used in this report came from NIMH Data Archive (NDA) Release 5.1 (DOI: 10.15154/z563-zd24). DOIs can be found at https://nda.nih.gov/abcd/abcd-annual-releases. The ABCD structural MRI data used in this report came from the fast-track data release. The raw data are available at https://nda.nih.gov/edit_collection.html?id=2573. Instructions on how to create an NDA study are available at https://nda.nih.gov/nda/tutorials/creating_an_nda_study.

## Code availability

Codes used in this manuscript is available at https://github.com/CNG-LAB/hippo_adversity.

Code and tutorials for HippUnfold preprocessing can be found at https://github.com/jordandekraker/hippunfold_toolbox.

## Acknowledgments

MM was supported by the Jacobs Foundation Research Fellowship through SLV. DYE was supported by the Max Planck Society through SLV. BS was supported by the German Federal Ministry of Education and Research (BMBF) and the Max Planck Society through SLV. MDH was supported by the German Research Foundation (DFG; Walter Benjamin Fellowship GZ: HE 10277/1-1). JDK was supported by a Natural Sciences and Engineering Research Council of Canada - Post-doctoral Fellowship (NSERC-PDF). FH was supported by the European Union’s Horizon 2020 Research Infrastructures Grant EBRAIN-Health 101058516 through SBE. BCB was supported by the Canadian Institutes of Health Research (CIHR), National Science and Engineering Research Council of Canada (NSERC), BrainCanada, Healthy Brains and Healthy Lives, the Canada Research Chairs (CRC) program, and the Centre for Excellence in Epilepsy at the Montreal Neurological Institute. SBE was supported by the European Union’s Horizon 2020 research and innovation programme (grant agreements 945539 [HBP SGA3], 826421 [VBC], and 101058516), the DFG (SFB 1451 and IRTG 2150), and the National Institute of Health (NIH; R01 MH074457). SLV was supported by the Max Planck Society through the Lise Meitner Excellence Program, the Jacobs Foundation Research Fellowship, the Hector Research Career Development Award, and the European Research Council (ERC) Starting Grant.

