## Supplementary Material for "Distinct spatial associations of adversity with hippocampal macro- and microstructure in early adolescence"

**Table S1.** Values indicate marginal and conditional $R^2$ for the linear mixed-effects models, representing variance explained by fixed effects alone and by the full model (fixed and random effects), respectively.

| Model | Region | $R^2$ marginal | $R^2$ conditional |
|---|---|---|---|
| **Volume** | | | |
| | Sum | 0.40 | 0.69 |
| | Subiculum | 0.32 | 0.56 |
| | CA1 | 0.30 | 0.57 |
| | CA2 | 0.24 | 0.48 |
| | CA3 | 0.22 | 0.57 |
| | CA4 | 0.19 | 0.42 |
| | Dentate gyrus | 0.24 | 0.59 |
| | Stratum radiatum lacunosum | 0.37 | 0.66 |
| **Thickness** | | | |
| | Mean | 0.20 | 0.54 |
| | Subiculum | 0.18 | 0.42 |
| | CA1 | 0.17 | 0.41 |
| | CA2 | 0.17 | 0.41 |
| | CA3 | 0.14 | 0.44 |
| | CA4 | 0.07 | 0.40 |
| | Anterior | 0.13 | 0.46 |
| | Anterior central | 0.06 | 0.43 |
| | Body | 0.08 | 0.38 |
| | Posterior central | 0.17 | 0.42 |
| | Posterior | 0.16 | 0.29 |
| **Gyrification** | | | |
| | Mean | 0.32 | 0.60 |
| | Subiculum | 0.19 | 0.47 |
| | CA1 | 0.18 | 0.45 |
| | CA2 | 0.15 | 0.40 |
| | CA3 | 0.19 | 0.55 |
| | CA4 | 0.22 | 0.42 |
| | Anterior | 0.12 | 0.42 |
| | Anterior central | 0.29 | 0.51 |
| | Body | 0.24 | 0.52 |
| | Posterior central | 0.13 | 0.48 |
| | Posterior | 0.12 | 0.45 |
| **T1w/T2w ratio** | | | |
| | Mean | 0.03 | 0.05 |
| | Subiculum | 0.03 | 0.05 |
| | CA1 | 0.03 | 0.06 |
| | CA2 | 0.03 | 0.05 |
| | CA3 | 0.03 | 0.05 |
| | CA4 | 0.03 | 0.04 |
| | Anterior | 0.03 | NA |
| | Anterior central | 0.03 | 0.05 |
| | Body | 0.03 | 0.07 |
| | Posterior central | 0.03 | 0.06 |
| | Posterior | 0.03 | 0.04 |

*CA: Cornu Ammonis*

**Table S2.** Values indicate the percentage of predictor and outcome variance explained by the latent component, mean test-set $R^2$ across response variables, and permutation-based significance of the component. All PLS-R models were restricted to a single latent component ($N_{train} = 3674$, $N_{test} = 1589$).

| Model | Explained variance (X) | Explained variance (Y) | $R^2$ test (mean) | Permutation $p$ |
|---|---|---|---|---|
| **Volume** | | | | |
| Sum | 18.37 | 35.70 | 0.36 | 0.001 |
| Proximal-distal axis | 18.33 | 23.35 | 0.32 | 0.001 |
| **Thickness** | | | | |
| Mean | 17.52 | 18.01 | 0.15 | 0.001 |
| Proximal-distal axis | 18.27 | 12.75 | 0.20 | 0.001 |
| Anterior-posterior axis | 17.92 | 10.63 | 0.12 | 0.001 |
| **Gyrification** | | | | |
| Mean | 18.06 | 27.26 | 0.25 | 0.001 |
| Proximal-distal axis | 18.05 | 15.69 | 0.20 | 0.001 |
| Anterior-posterior axis | 18.11 | 15.18 | 0.10 | 0.001 |
| **T1w/T2w ratio** | | | | |
| Mean | 13.56 | 2.50 | 0.04 | 0.001 |
| Proximal-distal axis | 13.57 | 2.43 | 0.05 | 0.001 |
| Anterior-posterior axis | 13.69 | 2.39 | 0.04 | 0.001 |

**Table S3.** Ethnicity and race description of the caregivers that did not answer the household income item in the MRI-eligible sample ($N = 8,666$).

| Variable | Category | Income response type | |
|---|---|---|---|
| | | decline to answer | do not know |
| **Caregiver ethnicity** | | | |
| | hispanic | 73 (21.6%) | 106 (33.87%) |
| | non-hispanic | 259 (76.63%) | 203 (64.86%) |
| | refuse to answer | 4 (1.18%) | 0 (0%) |
| | do not know | 2 (0.59%) | 4 (1.28%) |
| **Caregiver race** | | | |
| | white | 191 (56.51%) | 153 (48.88%) |
| | black or african american | 81 (23.96%) | 97 (30.99%) |
| | american indian or native american | 12 (3.55%) | 16 (5.11%) |
| | alaska native | 0 (0%) | 0 (0%) |
| | native hawaiian | 0 (0%) | 0 (0%) |
| | guamanian | 0 (0%) | 0 (0%) |
| | samoan | 1 (0.3%) | 1 (0.32%) |
| | other pacific islander | 1 (0.3%) | 0 (0%) |
| | asian indian | 9 (2.66%) | 1 (0.32%) |
| | chinese | 4 (1.18%) | 3 (0.96%) |
| | filipino | 4 (1.18%) | 4 (1.28%) |
| | japanese | 5 (1.48%) | 0 (0%) |
| | korean | 2 (0.59%) | 1 (0.32%) |
| | vietnamese | 2 (0.59%) | 1 (0.32%) |
| | other asian | 0 (0%) | 4 (1.28%) |
| | other race | 34 (10.06%) | 42 (13.42%) |
| | refuse to answer | 5 (1.48%) | 6 (1.92%) |
| | do not know | 9 (2.66%) | 13 (4.15%) |

**Table S4.** Caregivers' ethnicity and race distribution remain broadly comparable between MRI-eligible ($N = 8,666$) and final samples ($N = 5,263$), indicating that our selection procedures did not substantially introduce additional selection bias beyond that present in the ABCD dataset.

| Variable | Category | MRI-eligible sample | Final sample |
|---|---|---|---|
| **Ethnicity** | | | |
| | hispanic | 1325 (16.11%) | 795 (15.11%) |
| | non-hispanic | 6849 (83.29%) | 4443 (84.42%) |
| | refuse to answer | 16 (0.19%) | 7 (0.13%) |
| | do not know | 32 (0.39%) | 18 (0.34%) |
| | NA | 1 (0.01%) | 0 (0%) |
| **Race** | | | |
| | white | 6256 (76.08%) | 4142 (78.7%) |
| | black or african american | 1224 (14.89%) | 710 (13.49%) |
| | american indian or native american | 199 (2.42%) | 123 (2.34%) |
| | alaska native | 1 (0.01%) | 1 (0.02%) |
| | native hawaiian | 10 (0.12%) | 7 (0.13%) |
| | guamanian | 1 (0.01%) | 1 (0.02%) |
| | samoan | 6 (0.07%) | 4 (0.08%) |
| | other pacific islander | 19 (0.23%) | 15 (0.29%) |
| | asian indian | 57 (0.69%) | 32 (0.61%) |
| | chinese | 92 (1.12%) | 65 (1.24%) |
| | filipino | 77 (0.94%) | 43 (0.82%) |
| | japanese | 40 (0.49%) | 26 (0.49%) |
| | korean | 47 (0.57%) | 35 (0.67%) |
| | vietnamese | 23 (0.28%) | 13 (0.25%) |
| | other asian | 43 (0.52%) | 31 (0.59%) |
| | other race | 436 (5.3%) | 242 (4.6%) |
| | refuse to answer | 45 (0.55%) | 19 (0.36%) |
| | do not know | 92 (1.12%) | 43 (0.82%) |

**Table S5.** Agreement analysis between caregiver- (C) and youth-reported (Y) responses on the Family Environment Scale (Conflict Scale) across all respondents ($N = 11,832$), assessed using Cohen's kappa.

| Question | No (Y) | Yes (Y) | No (C) | Yes (C) | Agree (Chance) | Kappa | CI low | CI high |
|---|---|---|---|---|---|---|---|---|
| We fight a lot in our family | 8949 | 2883 | 9872 | 1960 | 0.67 | 0.12 | 0.10 | 0.15 |
| Family members rarely become openly angry | 7892 | 3940 | 5873 | 5959 | 0.50 | 0.06 | 0.04 | 0.08 |
| Family members sometimes get so angry they throw things | 10366 | 1466 | 10316 | 1516 | 0.78 | 0.12 | 0.08 | 0.15 |
| Family members hardly ever lose their tempers | 7524 | 4308 | 5705 | 6127 | 0.50 | 0.09 | 0.07 | 0.11 |
| Family members often criticize each other | 9529 | 2303 | 7960 | 3872 | 0.61 | 0.09 | 0.06 | 0.11 |
| Family members sometimes hit each other | 9220 | 2612 | 10383 | 1449 | 0.71 | 0.14 | 0.11 | 0.16 |
| If there's a disagreement in our family, we try hard to smooth things over and keep the peace | 10718 | 1114 | 10525 | 1307 | 0.82 | 0.01 | -0.03 | 0.05 |
| Family members often try to one-up outdo each other | 9133 | 2699 | 9773 | 2059 | 0.68 | 0.09 | 0.06 | 0.11 |
| In our family, we believe you don't ever get anywhere by raising your voice | 8954 | 2878 | 6065 | 5767 | 0.51 | 0.03 | 0.01 | 0.05 |

**Table S6.** Agreement analysis between caregiver- (C) and youth-reported (Y) responses on the Family Environment Scale (Conflict Scale) across the final sample ($N = 5,263$), assessed using Cohen's kappa.

| Question | No (Y) | Yes (Y) | No (C) | Yes (C) | Agree (Chance) | Kappa | CI low | CI high |
|---|---|---|---|---|---|---|---|---|
| We fight a lot in our family | 3977 | 1286 | 4380 | 883 | 0.67 | 0.12 | 0.09 | 0.16 |
| Family members rarely become openly angry | 3540 | 1723 | 2653 | 2610 | 0.50 | 0.07 | 0.04 | 0.10 |
| Family members sometimes get so angry they throw things | 4635 | 628 | 4575 | 688 | 0.78 | 0.11 | 0.06 | 0.16 |
| Family members hardly ever lose their tempers | 3343 | 1920 | 2579 | 2684 | 0.50 | 0.10 | 0.07 | 0.13 |
| Family members often criticize each other | 4227 | 1036 | 3573 | 1690 | 0.61 | 0.09 | 0.06 | 0.12 |
| Family members sometimes hit each other | 4062 | 1201 | 4607 | 656 | 0.70 | 0.15 | 0.11 | 0.19 |
| If there's a disagreement in our family, we try hard to smooth things over and keep the peace | 4803 | 460 | 4727 | 536 | 0.83 | -0.01 | -0.07 | 0.05 |
| Family members often try to one-up outdo each other | 4052 | 1211 | 4404 | 859 | 0.68 | 0.08 | 0.05 | 0.12 |
| In our family, we believe you don't ever get anywhere by raising your voice | 4038 | 1225 | 2623 | 2640 | 0.50 | 0.03 | 0 | 0.05 |

**Table S7.** Questionnaire items included in the Exploratory Factor Analysis, together with the Kiddie Schedule for Affective Disorders and Schizophrenia - Post-Traumatic Stress Disorder Module (KSADS-PTSD) items used for the inclusion criteria.

| Item | Question | Item | Question | Item | Question |
|---|---|---|---|---|---|
| crpbi_parent1_y | First caregiver makes me feel better after talking over my worries with him/her | parent_monitor_q1_y | How often do your guardians know where you are? | fam_enviro1_p/fes_youth_q1 | We fight a lot in our family |
| crpbi_parent2_y | First caregiver smiles at me very often | parent_monitor_q2_y | How often do your guardians know who you are with when you are not at school and away from home? | fam_enviro2_p/fes_youth_q2 | Family members rarely become openly angry |
| crpbi_parent3_y | First caregiver is able to make me feel better when I am upset | parent_monitor_q3_y | If you are at home when your guardians are not, how often do you know how to get in touch with them? | fam_enviro3_p/fes_youth_q3 | Family members sometimes get so angry they throw things |
| crpbi_parent4_y | First caregiver believes in showing his/her love for me | parent_monitor_q4_y | How often do you talk to your guardian about your plans for the coming day, such as your plans about what will happen at school or what you are going to do with friends? | fam_enviro4_p/fes_youth_q4 | Family members hardly ever lose their tempers |
| crpbi_parent5_y | First caregiver is easy to talk to | parent_monitor_q5_y | In an average week, how many times do you and your guardians, eat dinner together? | fam_enviro5_p/fes_youth_q5 | Family members often criticize each other |
| | | | | fam_enviro6_p/fes_youth_q6 | Family members sometimes hit each other |
| | | | | fam_enviro7_p/fes_youth_q7 | If there's a disagreement in our family, we try hard to smooth things over and keep the peace |
| | | | | fam_enviro8_p/fes_youth_q8 | Family members often try to one-up or outdo each other |
| | | | | fam_enviro9_p/fes_youth_q9 | In our family, we believe you don't ever get anywhere by raising your voice |
| **KSADS Item** | **KSADS Question** | **KSADS Item** | **KSADS Question** | **KSADS Item** | **KSADS Question** |
| ksads_ptsd_raw_761_p | Shot, stabbed, or beaten brutally by a non-family member | ksads_ptsd_raw_764_p | A non-family member threatened to kill your child | ksads_ptsd_raw_768_p | An adult outside your family touched your child in their privates, had your child touch their privates or did other sexual things to your child |
| ksads_ptsd_raw_762_p | Shot, stabbed, or beaten brutally by a grown-up in the home | ksads_ptsd_raw_765_p | A family member threatened to kill your child | ksads_ptsd_raw_769_p | A peer forced your child to do something sexually |
| ksads_ptsd_raw_763_p | Beaten to the point of having bruises by a grown-up in the home | ksads_ptsd_raw_767_p | A grown-up in the home touched your child in their privates, had your child touch their privates, or did other sexual things to your child | | |

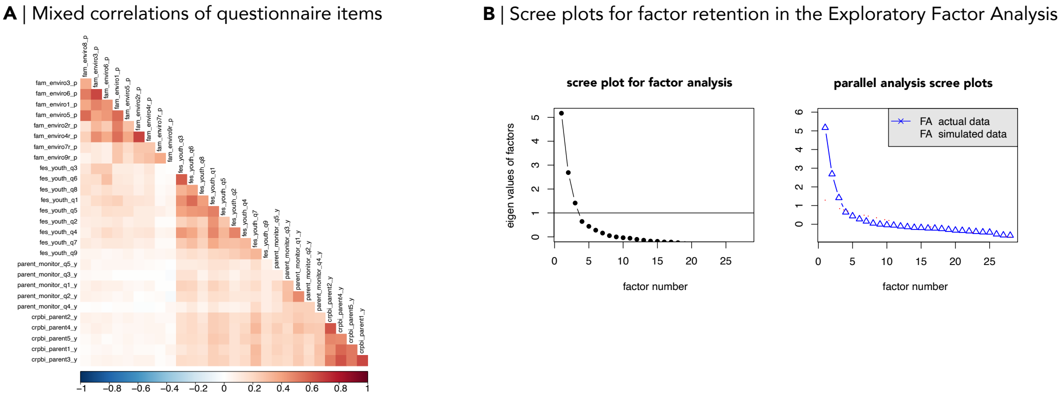


**Figure S1. Steps performed in the Exploratory Factor Analysis.** **A** | Kendall-based correlation matrix of ordinal and dichotomous questionnaire items included in the Exploratory Factor Analysis (EFA). **B** | Scree plots and parallel analysis supported the retention of three factors.

**Table S8.** Exploratory factor analysis loadings and internal consistency estimates for the final three-factor exploratory factor analysis solution, after removing loadings below 0.32 (∼10% of shared variance) during model refinement. One retained item (parent_monitor_q1_y) had a final loading of 0.302 after re-estimation of the final factor solution.

| Factor | Items | Loadings | Cronbach's $\alpha$ |
|---|---|---|---|
| Adverse family environment (youth) | | | 0.68 |
| | fes_youth_q1 | 0.757 | 0.62 |
| | fes_youth_q2 | 0.479 | 0.66 |
| | fes_youth_q3 | 0.703 | 0.65 |
| | fes_youth_q4 | 0.596 | 0.64 |
| | fes_youth_q5 | 0.594 | 0.65 |
| | fes_youth_q6 | 0.784 | 0.64 |
| | fes_youth_q7 | 0.345 | 0.68 |
| | fes_youth_q8 | 0.543 | 0.66 |
| Adverse family environment (caregiver) | | | 0.66 |
| | fam_enviro1_p | 0.782 | 0.62 |
| | fam_enviro2_p | 0.619 | 0.63 |
| | fam_enviro3_p | 0.622 | 0.64 |
| | fam_enviro4_p | 0.733 | 0.61 |
| | fam_enviro5_p | 0.643 | 0.62 |
| | fam_enviro6_p | 0.569 | 0.64 |
| | fam_enviro7_p | 0.365 | 0.66 |
| | fam_enviro8_p | 0.488 | 0.65 |
| | fam_enviro9_p | 0.351 | 0.67 |
| Neglectful parenting | | | 0.68 |
| | crpbi_parent1_y | 0.785 | 0.61 |
| | crpbi_parent2_y | 0.673 | 0.63 |
| | crpbi_parent3_y | 0.820 | 0.61 |
| | crpbi_parent4_y | 0.805 | 0.65 |
| | crpbi_parent5_y | 0.650 | 0.62 |
| | parent_monitor_q1_y | 0.302 | 0.68 |
| | parent_monitor_q4_y | 0.373 | 0.69 |

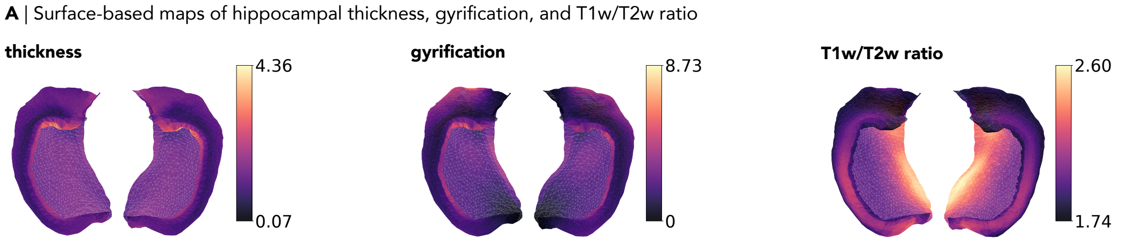


**Figure S2. Non-harmonized surface-based of hippocampal macro- and microstructural maps**. **A** | Surface-based maps show average non-harmonized feature values across participants retained in the final sample ($N = 5,263$).

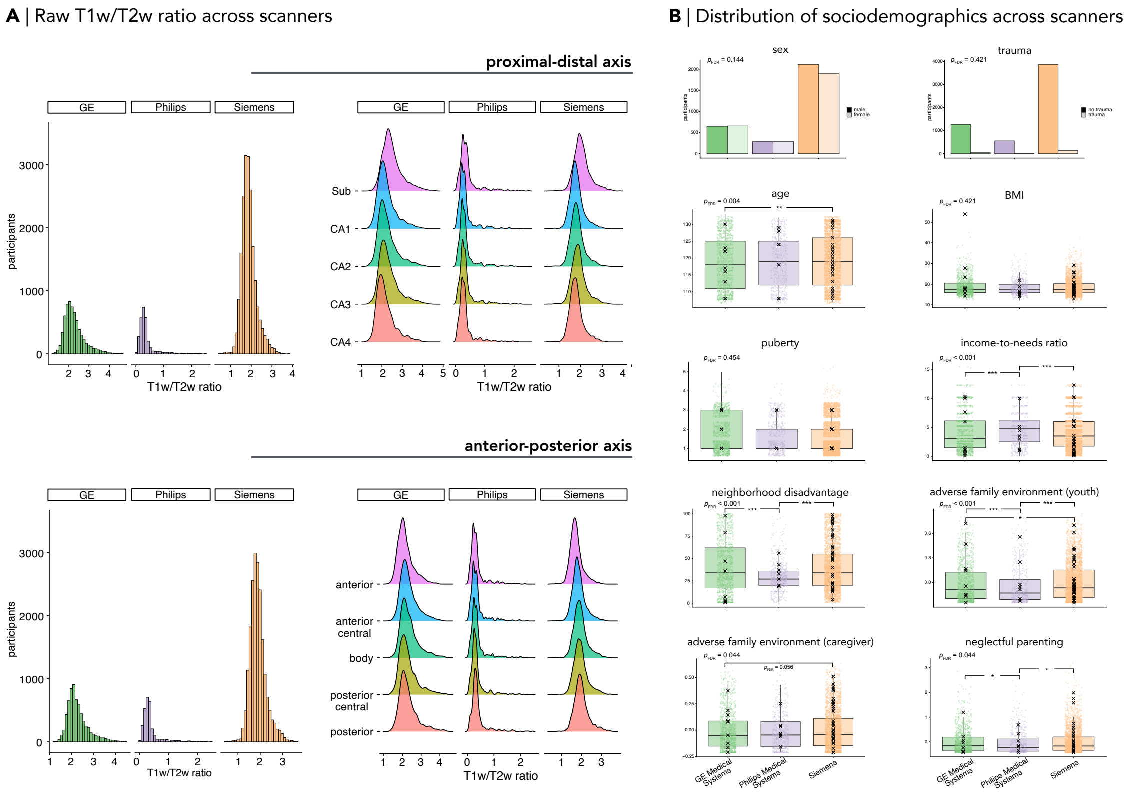


**Figure S3. Distribution of T1w/T2w ratios and sociodemographical variables across ABCD scanner manufacturers**. **A** | Participants scanned with Philips Medical Systems scanners exhibited systematically higher T1w/T2w ratio values compared to General Electric and Siemens scanners. Non-harmonized values are shown after removal of implausible ratios ($N = 1$) and 4$\pm$ standard deviation outliers ($N_{\text{GE}} = 9$, $N_{\text{Philips}} = 7$, and $N_{\text{Siemens}} = 36$). **B** | Participants scanned with Philips Medical Systems scanners also differed significantly in some biological and sociodemographic variables. Black crosses indicate the removed 4$\pm$ standard deviation outliers removed in subsequent analyses. *Sub: subiculum. CA: Cornu Ammonis.*
